# Easy to use and low cost leaf disease quantification workflow using Ilastik

**DOI:** 10.64898/2026.05.14.719059

**Authors:** Axel Prouvost, Léna Connesson, Titouan Le Gourrierec, Hélène Fréville, Jacques David, Baptiste Magnier, Clément Plessis

## Abstract

Accurate and reproducible assessment of foliar disease severity is essential for evaluating the performance of heterogeneous plant communities and understanding host-pathogen interactions. However, traditional visual scoring methods remain subjective, with limited precision, and difficult to scale in large phenotyping experiments. Here, we present a semi-automated image analysis workflow designed to quantify multiple foliar disease symptoms simultaneously on wheat flag leaves sampled from varietal mixtures. The workflow combines three methodological components: (i) a standardized protocol for leaf sampling and imaging, (ii) supervised machine learning segmentation using Random Forest implemented in Ilastik to classify multiple symptoms (powdery mildew and yellow rust), and (iii) a graphical user interface facilitating pipeline deployment by non-specialist operators. To evaluate the influence of image representation on classification performance, four color spaces (RGB, HSV, HLS, LAB) were systematically compared. The approach was validated using images of durum wheat flag leaves collected from a field experiment assessing eight-way varietal mixtures under natural fungal pressure. Cross-validation against manually annotated images demonstrated high segmentation accuracy across all symptom. Comparison among color spaces revealed only minor differences in performance. Overall, this workflow offers a cost-effective, annotation-efficient and reproducible alternative to deep learning approaches, leveraging open-source and actively maintained tools while requiring limited training data and enabling objective, reproducible and scalable disease phenotyping.

## 1 Introduction

In all ecosystems, plant communities are shaped by a complex network of biotic interactions. Among these, pathogens are key drivers of ecological and evolutionary dynamics, as they can significantly alter plant fitness depending on host susceptibility, environmental conditions, and pathogen virulence. In agrosystems, plant pathogens are a major cause of yield losses in many crops, accounting for up to 20–40% of potential production (Oerke, 2006). Among these, foliar diseases are the most widespread, affecting both the quality and quantity of agricultural production (Savary et al., 2019). Caused by fungi, bacteria, or viruses, these diseases typically manifest as visible symptoms on leaves. The qualitative and quantitative evaluation of such symptoms is therefore central to multiple disciplines, including plant pathology, epidemiology, ecology and genetics, and underpins applied activities such as epidemiological surveillance and cultivar resistance screening in agriculture. Visual assessment of disease symptoms remains relatively straightforward in ecosystems or agrosystems with low diversity, where species and genotypes are few and easily identifiable. In more complex systems, however, the co-occurrence of numerous species or genotypes complicates accurate visual scoring (Finckh et al., 2000; Cowger and Mundt, 2002; Bock et al., 2010).

In agriculture, crop diversification through multi-species cropping systems and varietal mixtures is increasingly promoted (Barot et al., 2017; Borg et al., 2018), leading to more complex systems for disease evaluation. For instance, genotypes with contrasting architectures (plant height, canopy density, etc.) can coexist within the same plot, creating a structurally heterogeneous canopy (Garrett and Mundt, 1999; Vidal et al., 2017). This physical heterogeneity locally modifies microclimate, rain splash interception, and short-distance spore dispersal, so that neighboring plants differing in susceptibility may experience contrasting pathogen pressures (Vidal et al., 2018). As a result, disease severity can vary substantially within a single field. To capture this variability, field assessments of foliar diseases typically rely on sampling large numbers of individual plant organs across locations and quantifying either the proportion of units affected, i.e., incidence, or the fraction of tissue showing lesions, i.e., severity (Madden and Hughes, 1999; mad, 2017; Vidal et al., 2020). In practice, however, the combination of large experimental designs and intensive sampling requirements makes visual scoring of each collected unit time-consuming, labor-intensive, and prone to inter-rater variability (Bock et al., 2010; mad, 2017). Computer vision approaches offer a promising avenue to overcome this bottleneck by enabling rapid, standardized severity assessments compatible with the sampling effort required in heterogeneous canopies.

Computer vision relies on three core tasks corresponding to increasing levels of spatial precision within an image. Classification assigns a label to an entire image or to a specific object within it; detection identifies and localizes objects by drawing bounding boxes around them; and segmentation partitions an image into distinct regions. Detection and segmentation both rely on classification, applied respectively at the object and region levels. Semantic segmentation represents the finest level of granularity, assigning a class label to every pixel in the image. Among the computational approaches available for these tasks, deep learning models based on neural networks —particularly convolutional neural networks— have become the backbone of countless predictive applications (LeCun et al., 2015). While these models provide powerful tools for plant disease classification and lesion segmentation (Mohanty et al., 2016; Rico-Fernández et al., 2019), their implementation requires programming skills, large annotated datasets and substantial computational resources for training (O’Mahony et al., 2020). Simpler machine learning approaches based on random forests offer an accessible alternative and have already proven effective for plant segmentation and pixel classification (O’Mahony et al., 2020). These models are computationally lightweight and can be implemented through user-friendly interfaces (Berg et al., 2019; Schindelin et al., 2012; Stritt et al., 2020). However, despite their accessibility, these tools still lack well-documented, end-to-end and flexible workflows for processing batch leaf images and quantifying multiple disease symptoms.

Image segmentation models are predominantly developed using the Red Green Blue (RGB) color space (Garcia-Garcia et al., 2018). Although RGB is the standard representation for digital images, alternative color spaces such as HSV (Hue, Saturation, Value), HLS (Hue, Lightness, Saturation), and LAB (Lightness, A, B; also called CIELAB) may enhance segmentation performance by isolating relevant visual features and improving discrimination ability (Cheng et al., 2001; Busin et al., 2009). The benefits of color space transformations have been demonstrated in several cases, particularly for distinguishing vegetation from soil (Hernández-Hernández et al., 2016). For instance, the HSV color space separates chromatic information (hue and saturation) from intensity (value), which can improve robustness to lighting variability in plant phenotyping and disease detection (Camargo and Smith, 2009). Similarly, the HLS representation incorporates a lightness component that more closely reflects perceived brightness (Kingdom, 2011), facilitating the detection of subtle color variations and improving contrast in shaded leaf areas(Johnson et al., 2021). The LAB color space was designed to approximate human color perception by ensuring that distances between colors correspond to perceived differences. As a result, subtle color shifts —such as the early yellowing or browning associated with leaf infection— can be more reliably detected and quantified, making LAB particularly suitable for characterizing and quantifying disease symptoms (Johnson et al., 2021). Despite these advantages, systematic evaluations of color space transformations for leaf disease segmentation remain scarce (Rico-Fernández et al., 2019; Canales et al., 2024).

In this study, we developed an accessible image analysis workflow to quantify foliar disease symptoms in complex systems. Leaf samples were collected from a field experiment in which 8-way varietal mixtures of durum wheat were exposed to natural epidemics of powdery mildew and rust. The workflow relies on Ilastik (Berg et al., 2019), a user-friendly, interactive machine learning tool developed for biologists that performs pixel classification with limited training data. Building on this tool, we implemented a fully automated processing pipeline integrating three main components: (i) a standardized protocol for acquiring high-resolution images of field-collected leaves using a portable flatbed scanner; (ii) an automated procedure to isolate individual leaves and convert images into multiple color spaces prior to classification, and (iii) a graphical user interface allowing users without programming expertise to process large batches of images, from raw scans to disease severity estimates. To evaluate the workflow, we generated training and test datasets from field-collected leaf samples using four color spaces and varying numbers of training images with annotated leaves, to determine the minimal annotation effort required for reliable segmentation. The best-performing model configurations were then applied to the full field dataset to assess which combinations of color space and training set size produced disease severity estimates most strongly correlated with mixture grain yield.

## 2 Material and Methods

### 2.1 Field experiment

We set up the experiment at Mauguio, southern France (INRAE– UE DIASCOPE – 43°36’49.8”N 3°58’49.5”E) on October 11^th^, 2022. The experiment consisted of 300 unique durum wheat varietal mixtures, each composed of eight genotypes sampled from a pool of 100 inbred lines derived from an Evolutionary Pre-breeding Population (David et al., 2014). We randomized the genotype position within plots by mixing 450 seeds per genotype to obtain seed pools sown in plots of dimensions of 1.36m width by 5m length. As we designed this experiment to test the effect of varietal mixtures on fungal diseases, we allocated the 300 plots into two blocks of 150 plots. The first block received a broad-spectrum fungicide treatment, while the second block remained untreated. On May 9^th^, 2023, we detected the presence of three diseases: powdery mildew, yellow rust and brown rust. On the same day, we randomly sampled 40 flag leaves per plot in the untreated block (eight batches of five leaves using a simple random sampling method as described by (Delp, 1986) from the 150 plots, for a total of 6000 leaves (see Figure .1.a step 1). On July 10^th^, 2023, we harvested all plots and measured their grain yield.

**Figure .1.**
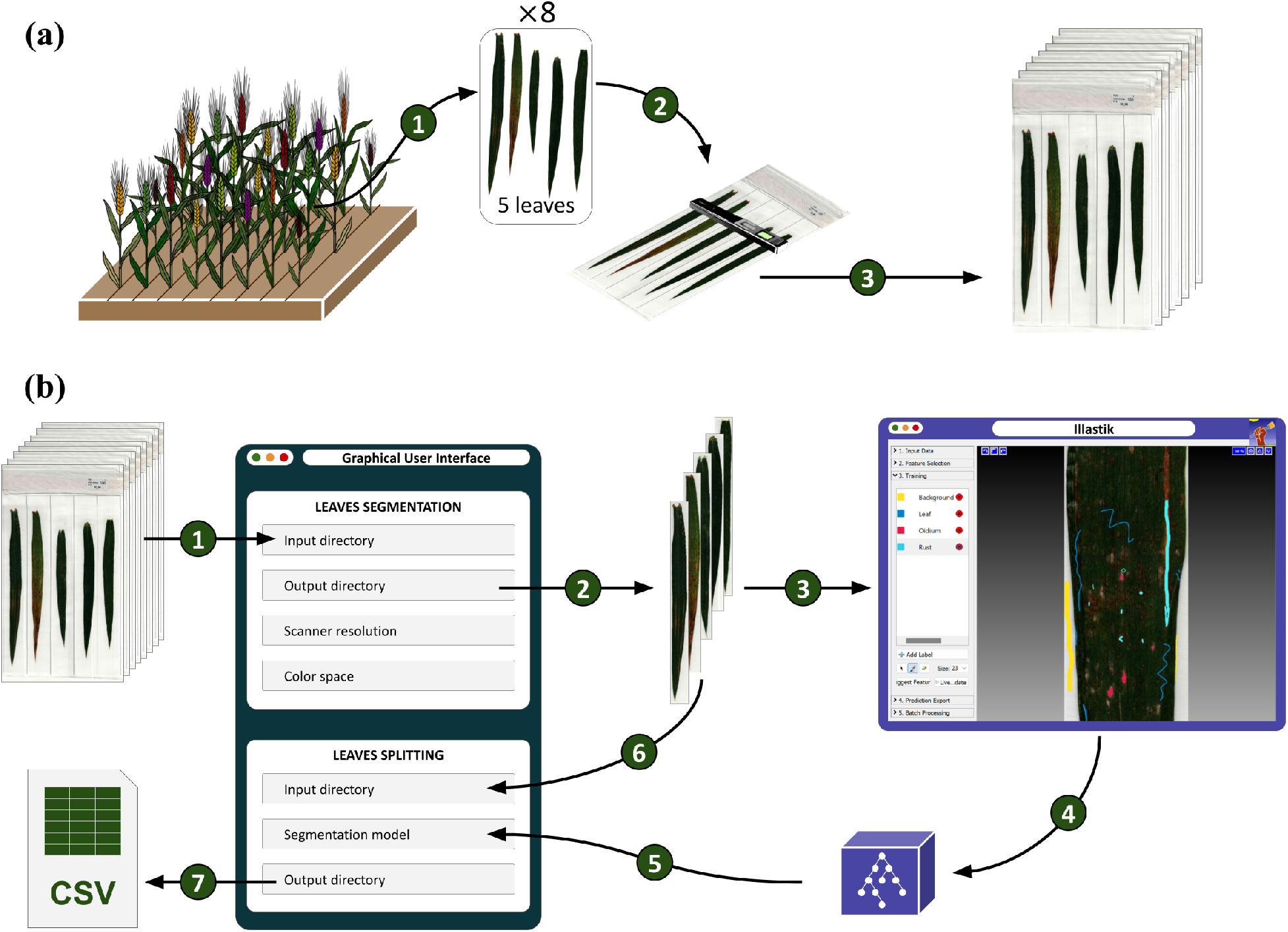
Workflow for leaf disease quantification. **a)** Image acquisition: step 1, randomly sample leaves; step 2, place the leaves on a sheet and cover them with a plastic film; step 3, scan the sheet with a portable scanner. **b)** Data processing: step 1, import sheet images in SegLeaf; step 2, select options and process to isolate each leaf, and choose a color space; step 3, use a subset of leaves to annotate with Illastik; step 4, train the model and export it; step 5, import your model in SegLeaf; step 6, import your dataset of isolated leaves to SegLeaf; step 7, run the segmentation to obtain a CSV report.

### 2.2 Leaf image acquisition

We arranged the leaves in batches of five on an A4 sheet corresponding to the dimensions of our portable scanners, in this case the IRIScan™ Book 5. We then covered the samples with plastic film to hold them in place during scanning (Figure .1.a step 2). After covering the samples with plastic film, we scanned the sheets at a resolution of 1200 dpi (Figure .1.a step 3). To isolate the scanned leaves on a uniform white background, we implemented a semi-automated segmentation protocol (Figure .1.b step 1-2). First, the RGB image was converted to greyscale and subjected to Gaussian blurring (5×5 kernel, *σ* = 1) in order to smooth out irregularities in the background (such as shadows or scanner residue) without altering the leaf contours. We then applied global thresholding using Otsu’s method (Otsu, 1979) to separate the pixels belonging to the leaves from those of the background. Closed contours were detected using OpenCV Python library (Bradski, 2000), then encapsulated in rectangular bounding boxes. This approach allowed for reliable leaf extraction (with > 95% accuracy on our dataset), even in the presence of minor variations in lighting or scan quality.

### 2.3 Color space transformation

Based on color spaces previously used in plant segmentation studies (Johnson et al., 2021), we selected four (Figure .2): RGB (Red, Green, Blue), HSV (Hue, Saturation, Value), HLS (Hue, Lightness, Saturation), and LAB (Lightness, A, B). To evaluate the impact of color representation on model performance, leaf isolation and color space transformation were performed separately for each representation using our graphical user interface, thereby, generating a dedicated training dataset for each color space (Figure .1.b step 2).

**Figure .2.**
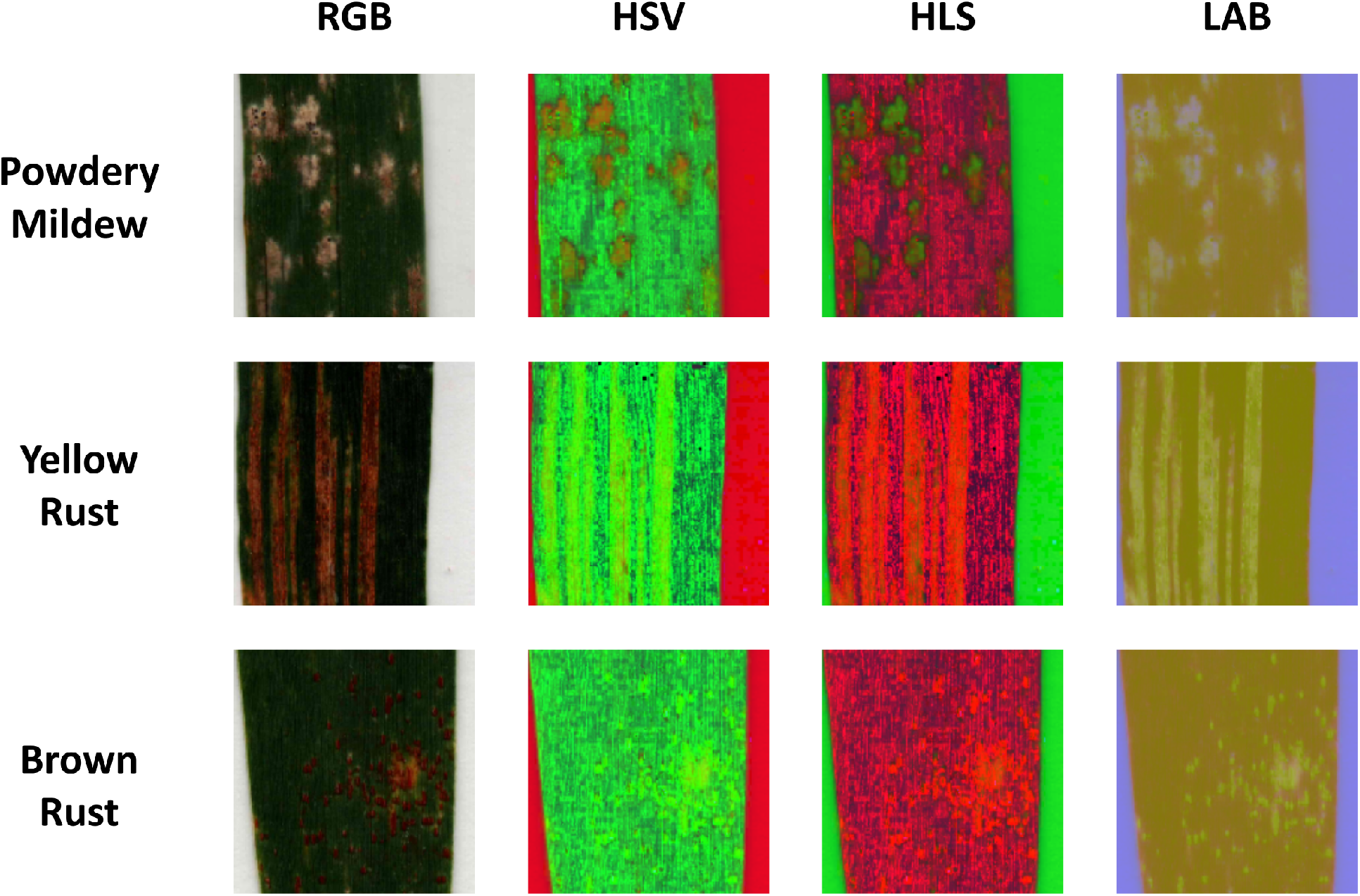
Disease symptoms observed on wheat leaves (powdery mildew, yellow rust and brown rust), for each color representation: RGB (Red, Green, Blue), HSV (Hue, Saturation, Value), HLS (Hue, Lightness, Saturation) and LAB (Lightness, A, B).

### 2.4 Pixel annotation and classification

Leaf annotation for training was performed using the *Ilastik* 1.4.0 software (Berg et al., 2019) with manual labeling of four distinct classes: background, healthy leaf tissue, powdery mildew, and rust. Because brown rust (Puccinia triticina) symptoms were too scarce to build a reliable training and test dataset, yellow and brown rust symptoms were merged into a single class (rust). For pixel classification, we selected the random forest algorithm due to its computational efficiency and native implementation in *Ilastik* (Breiman, 2001). The random forest model constructs an ensemble of decision trees trained on random subsets of data and features, thereby mitigating overfitting. Predictions are derived via majority voting across all trees, formalized as:

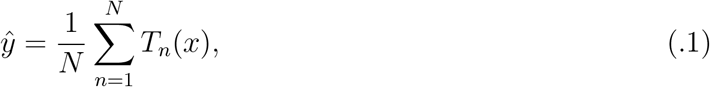

where *ŷ* is the predicted class label, *N* is the number of trees, and *T*_*n*_(*x*) is the prediction from the *n*-th tree for input *x*. Each tree consists of internal nodes and leaves: internal nodes apply decision rules to the input x to determine the path to subsequent nodes, terminating at a leaf node that provides the final prediction *T*_*n*_(*x*).

Ilastik integrates six fundamental filters that can be applied across multiple spatial scales defined by the *σ* (sigma) parameter, allowing the extraction of multi-scale features for image classification or segmentation. These filters capture patterns ranging from fine details (e.g., edges or local textures) to broader structures (e.g., cellular shapes or homogeneous regions). The choice of *σ* values is therefore critical for adapting the analysis to the resolution and characteristics of the images. In practice, *σ* values can be grouped into three ranges: small scales (*σ* ∈ [0.3, 1.0]), suited for detecting fine details such as small contours; intermediate scales (*σ* ∈ [1.6, 3.5]), capturing moderate structures such as pustules or medium-sized fibers; and large scales (*σ* ∈ [5, 10]), used to identify broader patterns such as tissue regions or large objects. Each filter can be applied at one or several of these scales, enabling hierarchical image feature extraction. Table .1 summarizes these filters, their categories, typical applications and the *σ* values used in our models.

**Table 1.**
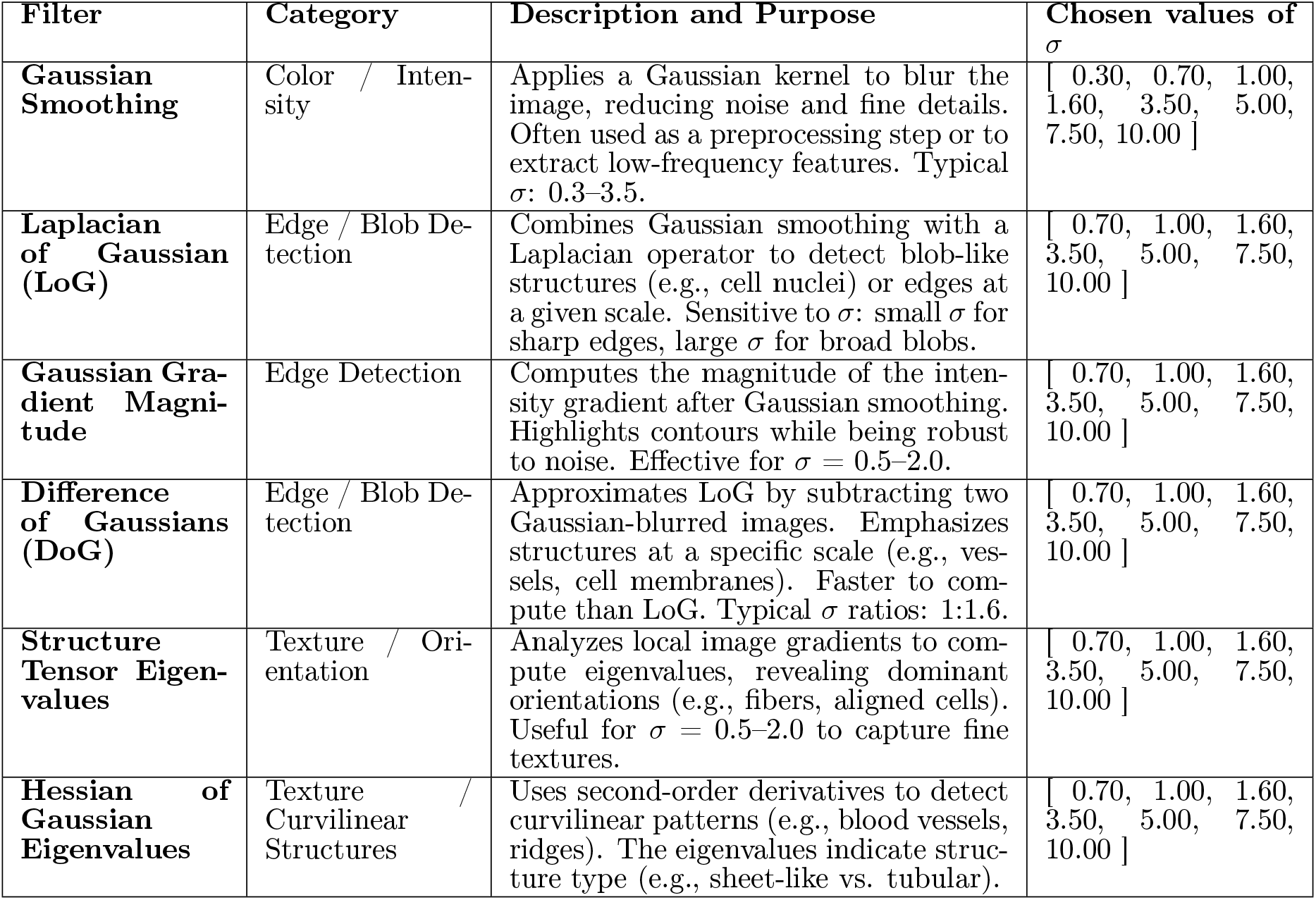
Summary of *Ilastik* feature filters, their typical applications and the *σ* parameters chosen for our classification models.

After defining the filter parameters (Table .1), we designed the training procedure to test two factors: training set size and color space representation. To assess the effect of training size, we annotated 15 leaf images in RGB and used these annotations to train models with subsets of 5, 10, and 15 leaves. To assess the effect of color space, the same annotated leaves were used to train additional models after transforming the images into HSV, HLS, and LAB representations. The combination of three training dataset sizes (5, 10, and 15 leaves) and four color spaces (RGB, HSV, LAB, and HLS) resulted in a total of twelve models.

### 2.5 Model evaluation

To evaluate the performance of our models, we created a validation dataset by manually annotating each of the four classes on 48 leaves using the VGG Image Annotator (VIA) 2.0.12 (Dutta and Zisserman, 2019). Using this dataset, we evaluated the performance of the models according to several complementary metrics ranging from 0 to 1 and based on the classification of true positive (*TP*), false positive (*FP*) and false negative (*FN*) pixels respectively:

The *Precision* criterion reflects the reliability of the model’s predictions by measuring the proportion of predicted positive pixels that are correctly classified, with high values indicating fewer false detections.

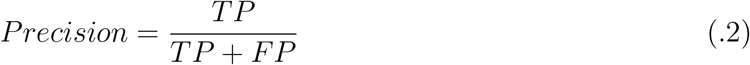

The *Recall* criterion measures completeness by quantifying how much of the true diseased region is successfully recovered by the model, with high values indicating a lower rate of false negatives.

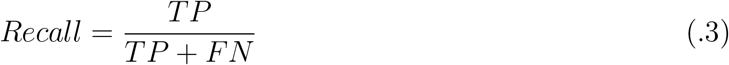

Since a model may achieve high precision at the expense of recall (*and vice versa*), we computed the *F*_1_ score by combining both metrics through their harmonic mean as follows:

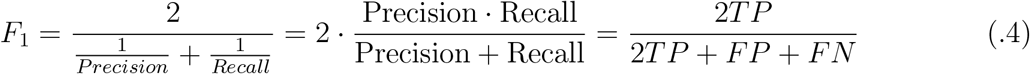

High values reflect a better trade-off between false positives and false negatives. This metric is thus particularly useful when looking for a single indicator that accounts for both false positives and false negatives.

The *IntersectionoverUnion*(*IoU*) criterion evaluates the spatial overlap between predicted and annotated regions, with higher values indicating greater agreement between segmentation outputs and ground truth. The *IoU* is especially valuable for assessing segmentation quality as it penalizes both over- and under-segmentation simultaneously.

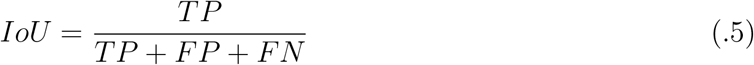

For each of the twelve models, we assessed its accuracy to predict the area of each segmentation class. To do this, we used the manual annotation of the 48 leaves composing the test dataset, hereafter referred to as “ground truth”. We then computed the squared Pearson correlation coefficient (*R*^2^) between the ground truth area and the corresponding predictions for each class.

Finally, assuming that fungal diseases reduce crop production, as documented in the literature, we tested the relationship between the model’s predictions of disease severity and observed mixture yields. For each color space, we first determined the optimal training dataset size (5, 10, or 15 leaves) based on the two complementary approaches mentioned above: the pixel classification performance metrics and the *R*^2^ values obtained for the estimated leaf areas affected by rust and powdery mildew. Using the four resulting models (one per color space), we then fitted linear regressions at the plot level, with disease severity as a predictor of yield. Three disease indicators were evaluated separately: rust severity, powdery mildew severity, and their combined severity (1−healthy leaf). These analyses were performed independently for each color space to assess whether automated segmentation could provide reliable predictions of yield loss. Finally, to test whether color space choice significantly affected predictive performance, we compared pairwise regression models using likelihood ratio tests (LRTs).

## 3 Results

### 3.1 Model performance

Since background and healthy leaf tissue covered substantially larger areas than diseases, we divided the presentation of the results into two groups: large area classes (Figure .3.a) and small area classes (Figure .3.b).

**Figure .3.**
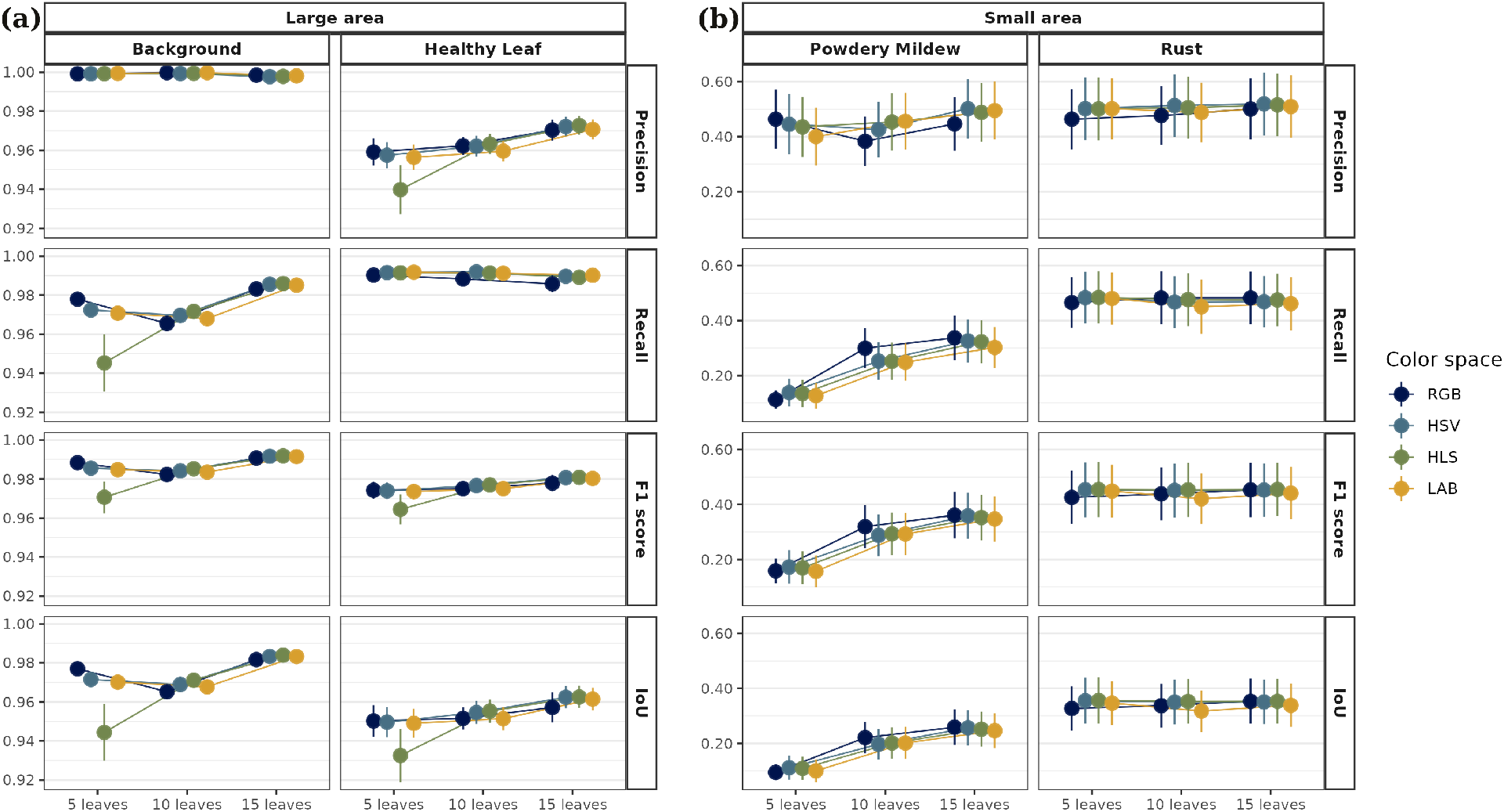
Comparison of pixel classification metrics for wheat leaf disease segmentation. Performance was evaluated across four color spaces (RGB, HSV, HLS, LAB) and three training set sizes (5, 10, 15 leaves) for **(a)** large area classes: Background and Healthy Leaf; and **(b)** small area classes: Powdery Mildew and Rust. Error bars indicate 95% confidence intervals (n = 48).

**Figure .4.**
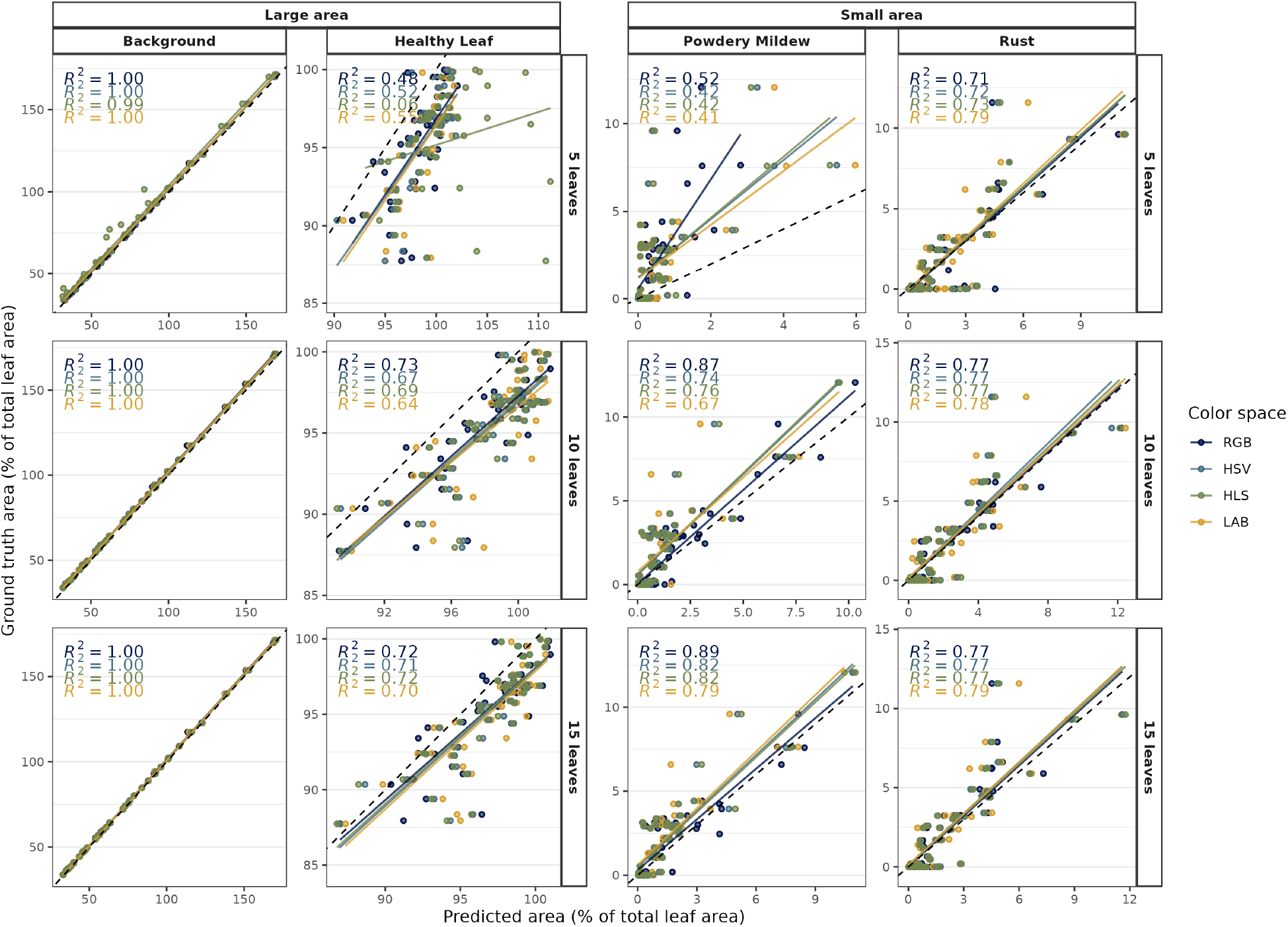
Linear regression of predicted versus ground truth area across four color spaces (RGB, HSV, HLS, LAB), three training set sizes (5, 10, and 15 leaves), and four segmentation classes, shown separately for **a)** large-area classes (Background and Healthy Leaf) and **b)** small-area classes (Powdery Mildew and Rust). R^2^ values are displayed for each combination.

**Figure .5.**
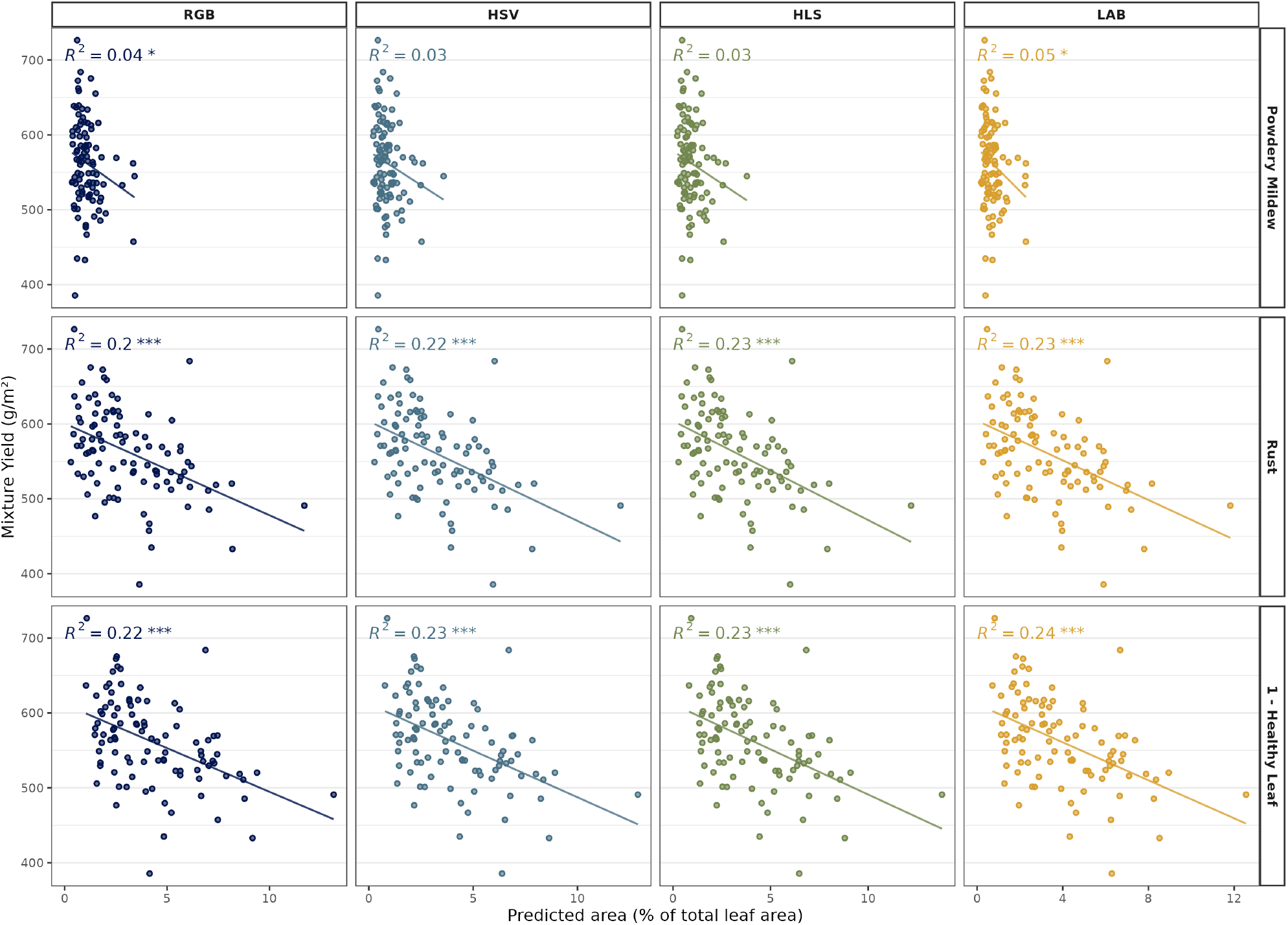
Relation between predicted disease severity by the best model and varietal mixture yield. Linear regressions were fitted using leaf area predictions for rust, powdery mildew, and their combined severity across four color spaces (RGB, HSV, HLS, LAB). Only results from the 15-leaf training model are presented. *R*^2^ values and regression *P* -values are displayed for each combination with the following significance levels: * *P* < 0.05, ** *P* < 0.01, *** *P* < 0.001

#### 3.1.1 Leaf and disease detections

The twelve segmentation models performed very well in distinguishing large area classes: background from leaf, and diseased pixels from healthy pixels (Figure .3.a). Indeed, both classes achieved high classification performance, with *Precision* ranging from 0.93 to 1, *Recall* from 0.93 to 1, *F*_1_ score from 0.96 to 1 and *IoU* from 0.92 to 1. Estimates of background and healthy tissue areas exhibited narrow confidence intervals for each metric (Figure .3.a). For both classes, the HLS color space consistently underperformed compared to the other color spaces. For the background class, a decrease in performance was observed across all color spaces when increasing the training set from 5 to 10 leaves; however, this gap was entirely recovered when the training set was further increased from 10 to 15 leaves. Additionally, increasing the number of training leaves from 5 to 15 generally improved model performance for detecting healthy leaf pixels.

#### 3.1.2 Classification is disease dependent

The twelve segmentation models were less performant in distinguishing small area classes, *i*.*e*., diseased pixels from healthy pixels for each disease (Figure .3.b). Estimates of powdery mildew and rust areas displayed substantially wider confidence intervals than large area estimates (Figure .3). In addition, no significant differences in performance were observed among color spaces across diseases and training set sizes (5, 10 or 15 leaves). For powdery mildew, *Precision* showed no improvement with increasing leaf number. In contrast, *Recall, F*_1_ score, and *IoU* increased significantly from 5 to 10 leaves, with only marginal gains from 10 to 15 leaves, leading to optimal performance with 15 leaves. For rust, detection performance was consistently higher than for powdery mildew across all metrics, except for *Precision*, which was comparable to that obtained for powdery mildew with 15 leaves. Furthermore, for all metrics, optimal performance was achieved with only 5 leaves across all color spaces, with no further improvement when increasing the training set size. Hence, the performance of the segmentation models revealed substantial differences between the two diseases.

#### 3.1.3 Accuracy of models in estimating the area of each class

Linear regression analyses revealed distinct performance patterns across the four segmentation classes depending on training dataset size. Background segmentation achieved perfect prediction accuracy (*R*^2^ = 1.00) across all color spaces and training set sizes. Prediction of healthy leaf area showed moderate performance (*R*^2^ = 0.06–0.73), but improved markedly when the training set increased from 5 to 10–15 training leaves, with all color spaces converging toward similar accuracy (*R*^2^ = 0.70–0.72). For powdery mildew, model performance strongly depended on training dataset size. With only 5 training leaves, *R*^2^ values remained low (0.41–0.52), but increased markedly with 15 leaves (0.79–0.89). In contrast, models of rust prediction exhibited more stable performance across configurations, with *R*^2^ values ranging from 0.71 to 0.79 regardless of training set size. Differences among color spaces were limited. For powdery mildew with 15 training leaves, RGB achieved the highest accuracy (*R*^2^ = 0.89), followed by HSV and HLS (*R*^2^ = 0.82) and LAB (*R*^2^ = 0.79). For rust quantification, LAB achieved the highest accuracy overall (*R*^2^ = 0.79).

### 3.2 Application to the full field dataset

Based on the comparative analysis of segmentation accuracy and disease area prediction, we retained models trained with 15 leaves, which provided the best overall performance across all configurations. In addition, all four color spaces were used for simultaneous disease detection and quantification, as their performance overall converged when the training set exceeded 10 leaves. The four resulting models were therefore applied to the full dataset using 40 leaves per plot. Due to the loss of data for 53 plots during image acquisition, subsequent yield regression analyses were performed on 97 mixtures. Predicted powdery mildew severity explained only a small proportion of yield variance, with *R*^2^ values ranging from 3% (HSV, *P* > 0.05) to 5% (LAB, *P* < 0.05), while HSV and HLS models did not reach statistical significance. In contrast, predicted rust severity explained a substantially larger proportion of yield variation across all color spaces, with *R*^2^ values ranging from 20% (RGB, *P* < 0.001) to 23% (LAB, *P* < 0.001). When combining both diseases into a single metric (1−healthy leaf), all color spaces significantly explained yield variation, with *R*^2^ values ranging from 22% (RGB, *P* < 0.001) to 24% (LAB, *P* < 0.001). Although the LAB color space consistently explained the largest proportion of variance across all disease metrics, likelihood ratio tests comparing models across color spaces revealed no significant differences.

## 4 Discussion

### 4.1 Overview of the proposed pipeline

Accurate phenotyping of foliar diseases remains challenging in complex ecosystems and agrosystems, particularly when several diseases need to be assessed simultaneously. Our study addresses this challenge by proposing a high-throughput phenotyping method that combines standardized leaf sampling with machine-learning-based image analysis to quantify the co-occurrence of rust and powdery mildew on leaves in durum wheat varietal mixtures, each composed of eight genotypes. Our user-friendly method relies on a Python package interfaced with the Ilastik annotation environment, leading to a comprehensive pipeline that incorporates color space transformations to enhance disease segmentation on high-resolution leaf images. It further combines image acquisition via a portable scanner, semi-automated leaf segmentation, and random-forest classification—enabling reliable estimation of multi-disease leaf infection.

Although pixel-level classification performance varied across classes, our results demonstrate that this flexible pipeline provides a robust solution specifically designed for multi-class segmentation.

### 4.2 Training dataset size

The impact of training set size on segmentation performance varied depending on the class considered. For the background class, which exhibits a relatively uniform color, performance reached its maximum with as few as 5 training leaves regardless of the color space used. For healthy leaf tissue, optimal performance was attained with 10 training leaves. For disease classes, the response to increasing training set size was more variable: rust classification performance rapidly reached a plateau, suggesting that fewer training samples are needed to capture the spectral characteristics of rust symptoms, whereas powdery mildew classification benefited from larger training sets, likely due to the greater variability in symptom appearance. Overall, configurations with 10 or 15 training leaves yielded the most robust area estimations, particularly for the disease classes. Although the number of leaves provides a practical proxy for the amount of annotated data, it is important to note that the total number of annotated pixels increases consistently with each additional leaf included in the training set. As a result, a small number of leaves exhibiting a broad range of symptom expression can be sufficient to train a reliable model, provided that the diversity of symptoms is adequately represented. From a practical standpoint, this minimal annotation requirement offers substantial time savings compared to conventional deep learning approaches, which require hundreds to thousands of annotated images to achieve similar segmentation performance (Barbedo, 2018). Moreover, the low training requirement facilitates rapid model retraining when encountering new symptom types or different species.

### 4.3 Color space choice

The four color spaces tested (RGB, HSV, HSL, and LAB) achieved comparable overall segmentation performance. This result contrasts with several studies that have reported superior performance of color spaces separating chromatic information from luminance (Velastegui and Pedersen, 2021; Johnson et al., 2021). Nevertheless, among the four color spaces tested, LAB was slightly more performant across most configurations in terms of standard classification metrics. Several studies have similarly reported that LAB-based classifiers provided more accurate estimations of lesion area for various foliar diseases (Velastegui and Pedersen, 2021; Johnson et al., 2021). However, this advantage did not consistently translate into better area estimation. For powdery mildew in particular, LAB-based models exhibited lower agreement between predicted and ground truth diseased area compared to RGB, with a *R*^2^ approximately 10% lower at 15 training leaves. For rust, no significant differences among color spaces were observed regardless of the evaluation method. These findings suggest that the advantages of color space selection are disease-dependent, and that caution is warranted when choosing a single color space for multi-disease models. Different pathogens may exhibit spectral characteristics that are best captured by different color transformations. Rather than applying a universal color space across all diseases, it may be more effective to develop disease-specific models, each optimized with the most appropriate color transformation. This hypothesis should be explored when designing classification models.

### 4.4 Performance across disease symptoms

The segmentation models exhibited markedly different performance levels between powdery mildew and rust. Powdery mildew detection was considerably more challenging, as reflected by lower F1 scores and wider confidence intervals across all tested configurations. Several factors can account for this reduced performance. The spectral similarity between powdery mildew symptoms, which manifest as whitish to pale grey patches on the leaf area, and the scanning background likely caused confusion between these two classes, leading to misclassification of background pixels as diseased tissue and vice versa. Background color has been identified as a critical factor in image-based disease assessment, as suboptimal choices can markedly reduce segmentation accuracy (Barbedo, 2014; Xiao et al., 2020). Furthermore, unlike the well-delineated yellow rust strips or brown rust pustules, powdery mildew infections exhibit irregular, discontinuous edges without clear geometric shapes, making precise manual annotation inherently challenging and contributing to model errors. In contrast, rust symptoms were consistently detected with higher accuracy across all configurations, achieving optimal performance with fewer training samples. This can be attributed to several complementary factors: their orange to brown coloration is spectrally distinct from both healthy leaf tissue and the background, enabling effective pixel classification even with limited training data, and their well-defined, roughly circular pustules arranged in strips with clear boundaries facilitate both automated detection and accurate manual annotation. These findings align with previous studies indicating that diseases producing visually contrasting lesions with sharp boundaries are more readily quantified through image analysis (Bock et al., 2010; Barbedo, 2016).

### 4.5 Workflow accessibility

In this study, we propose a solution accessible to researchers without computer science expertise for developing their own disease segmentation models. The use of Ilastik, a software with an intuitive graphical interface, requires no programming skills (Berg et al., 2019) and provides a long term solution as it is an open-source and maintained tool. Combined with our simple interface for image pre-processing and post-processing, it allows users to move from raw leaf scans to disease severity estimates. Accessible workflows for plant disease quantification remain scarce in the literature, as most existing pipelines or tools either require programming proficiency or rely on proprietary software. The accessibility and standardization of our approach also facilitate its extension to other applications beyond the pathosystems investigated here. The classifier algorithms within Ilastik can be trained to recognize any visually distinguishable features, including insect damage, nutritional deficiencies, senescence patterns, or phenological stages. Furthermore, the modular structure of the workflow —separating image acquisition, pre-processing, and classification— facilitates adaptation to new use cases: users only need to annotate the features relevant to their specific phenotyping targets within the Ilastik interface, after which the same workflow can be applied consistently.

### 4.6 Practical recommendations for implementation

Based on our results and the experience gained during the development and application of the pipeline, several methodological recommendations can be made for users seeking to implement or adapt this workflow. Although JPEG compression remains standard in many practical agricultural imaging applications (Arnal Barbedo, 2013), it introduces color-metric degradation through lossy encoding and chroma subsampling. To maximise segmentation accuracy, users should consider using scanners capable of exporting to lossless formats such as PNG or TIFF, which preserve full color information and improve the discrimination of subtle symptom variations. Finally, while Ilastik’s default parameters were retained throughout this study to simplify the methodology and ensure reproducibility, systematic optimization of the *σ* parameters governing the feature extraction filters could further improve classification performance for specific types of symptoms, representing a promising avenue for future applications of the pipeline.

## Acknowledgements

We thank the DIASCOPE platform (INRAE, Mauguio, France) for providing all the facilities and technical support in the field experiment; the Ge2pop Team (AGAP Institute, France) for their help in leaf sampling and scanning. This work was supported by the Agence Nationale de la Recherche (ANR) (project SCOOP, grant no. ANR-19-CE32-0011; and project MOBIDIV, grant no. ANR-20-PCPA-0006).

## Code and Data Availability

The method and associated scripts developed in this work are freely available to the research community. All code is hosted in a public GitHub repository at https://github.com/titouanlegourrierec/SegLeaf, which includes the full implementation of the method including the graphical interface and documentation to guide users through the analysis pipeline.

